# A polygenic *p* factor for major psychiatric disorders

**DOI:** 10.1101/287987

**Authors:** Saskia Selzam, Jonathan R. I. Coleman, Avshalom Caspi, Terrie E. Moffitt, Robert Plomin

**Author notes:** **Corresponding author**: Saskia Selzam.

## Abstract

It has recently been proposed that a single dimension, called the *p* factor, can capture a person’s liability to mental disorder. Relevant to the *p* hypothesis, recent genetic research has found surprisingly high genetic correlations between pairs of psychiatric disorders. Here, for the first time we compare genetic correlations from different methods and examine their support for a genetic *p* factor. We tested the hypothesis of a genetic *p* factor by using principal component analysis on matrices of genetic correlations between major psychiatric disorders estimated by three methods – family study, Genome-wide Complex Trait Analysis, and Linkage-Disequilibrium Score Regression – and on a matrix of polygenic score correlations constructed for each individual in a UK-representative sample of 7,026 unrelated individuals. All disorders loaded on a first unrotated principal component, which accounted for 57%, 43%, 34% and 19% of the variance respectively for each method. Our results showed that all four methods provided strong support for a genetic *p* factor that represents the pinnacle of the hierarchical genetic architecture of psychopathology.

## Introduction

High comorbidity rates among psychiatric disorders^1^ have led to research investigating higher-order dimensions for psychopathology, including Internalizing (e.g., anxiety and depression), Externalizing (e.g., hyperactivity and conduct disorder), and Psychotic Experiences (e.g., schizophrenia and bipolar disorder)^2^. However, these higher-order dimensions also correlate with each other^3^, which suggests the possible existence of a general factor of psychopathology^4^.

This general factor has been called the *p* factor^5^ because it parallels the *g* factor of general intelligence^6^ which sits on top of the widely accepted hierarchical model of cognitive abilities^7^. *g* accounts for 40% of the variance of diverse cognitive tests, is stable across the life course, and is one of the best predictors of educational, occupational and health outcomes^8^. Analogous to *g*, the *p* factor captures the shared variance across psychiatric symptoms, and predicts a multitude of poor outcomes and general life impairment^9,10^.

Family studies support the hypothesis of a genetic *p* factor in that genetic influences on psychopathology appear to be general across disorders rather than specific to each disorder. For example, psychiatric disorders do not breed true – parental psychopathology predicts offspring psychiatric disorders but with little specificity^11^. Family research has found substantial genetic correlations between pairs of disorders, such as Major Depression and Generalized Anxiety Disorder^12^ and Schizophrenia and Bipolar Disorder^13^. Genetic overlap between internalizing and externalizing higher-order constructs has also been noted^14^, consistent with the hypothesis of a general *p* factor. The culmination of this research is a recent study of more than three million full- and half-siblings using Swedish national register data that found evidence for a general genetic factor that pervades eight major psychiatric disorders as well as convictions for violent crimes^15^. Although genetic correlations were not presented, the average loading was 0.45 on a general genetic factor.

Genomic research also supports the hypothesis of a genetic *p* factor. The first hint came from genome-wide association (GWA) findings that single-nucleotide polymorphisms (SNPs) found to be associated with Schizophrenia were also associated with Bipolar Disorder^16^. Also in 2013, genetic correlations were first estimated from linear mixed model analyses (Genome-wide Complex Trait Analysis, GCTA) of individual genotype data for five psychiatric disorders in the Psychiatric Genomics Consortium (PGC)^17^. Schizophrenia, Bipolar Disorder and Major Depressive Disorder yielded the highest genetic intercorrelations (average = 0.53); the average genetic correlation among the five disorders, including Autistic Spectrum Disorder and Attention-Deficit/Hyperactivity Disorder, was 0.22.

Linkage-Disequilibrium Score Regression (LDSC)^18^ has made it possible to estimate genetic correlations from GWA summary statistics rather than requiring genotype data for individuals. This method is based on correlations in effect sizes across disorders taking into account linkage disequilibrium and the SNP heritabilities of the disorders. LDSC genetic correlations derived from summary GWA statistics for the same five PGC disorders are remarkably similar to the GCTA genetic correlations described above that used individual genotype data^19^. A recent LDSC analysis of eight psychiatric disorders again showed considerable correlations between Schizophrenia, Bipolar Disorder and Major Depressive Disorder (average = 0.41), and yielded an average genetic correlation of 0.21^20^, highlighting the relevance of testing the hypothesis of a genetic *p* factor.

Another approach that has not yet been systematically applied to test for a genetic *p* is to correlate genome-wide polygenic scores (GPS), although some GPS correlations between pairs of psychiatric disorders have been reported^21^. A GPS for a disorder is created for an individual by summing the alleles shown in GWA studies to be associated with the disorder, after weighting the alleles by the strength of their association^22^. The previously described PGC dataset was used to create polygenic scores for each of the five disorders^16^, and polygenic scores for Schizophrenia, Bipolar Disorder and Major Depressive Disorder predicted liability variance in the other disorders, again suggesting genetic overlap. However, as new GWA studies have been published since for Schizophrenia, Attention-Deficit/Hyperactivity Disorder and Autism Spectrum Disorder with considerably increased sample sizes, replication is needed. GPS correlations between disorders are related to genetic correlations, but differ from the genetic correlations estimated from other methods because they index both the relationship between individual-specific genetic effects for traits in the population and genetic effects derived from an independent analysis. Nonetheless, GPS correlations provide another opportunity to test the hypothesis of a genetic p factor.

Based on the overwhelming evidence that favours a general *p* factor, we test whether a general *p* factor also emerges from genetic data. In the present study, we bring together genetic correlations for major psychiatric disorders derived from four genetic methods (family, GCTA, LDSC, and GPS). We applied principal component analysis to correlation matrices derived from these four methods and estimate the amount of genetic variance explained by a genetic *p* factor. For the GPS approach, we constructed GPS for eight psychiatric disorders for each individual in a sample of 7,026 unrelated individuals from the Twins Early Development Study (TEDS)^23^.

Our hypotheses were that (i) a general genetic factor would emerge from factor analyses of correlations derived from each of the four genetic methods, and that (ii) this genetic *p* factor would account for a substantial portion of the variance of the correlation matrices.

## Results

### Genetic correlations

Figure 1 presents the genetic correlations from family analysis, GCTA and LDSC, and the correlations from GPS analysis. The average genetic correlations were 0.49 for family analysis, 0.24 for GCTA and 0.31 for LDSC, indicating general genetic overlap among psychiatric disorders. The average GPS correlation was lower (0.06), as expected. However, genetic correlations for all four genetic approaches clustered in a strikingly similar way. Most notably, the average genetic correlations between Schizophrenia, Bipolar and Depression were consistently the largest in magnitude – 0.67 for family analysis, 0.53 for GCTA, 0.59 for LDSC, and 0.21 for GPS. High genetic correlations were not driven by larger heritability estimates for these traits in comparison to the other disorders (see Supplementary Tables S2 and S3 for SNP-h^2^ estimates).

**Figure 1.**
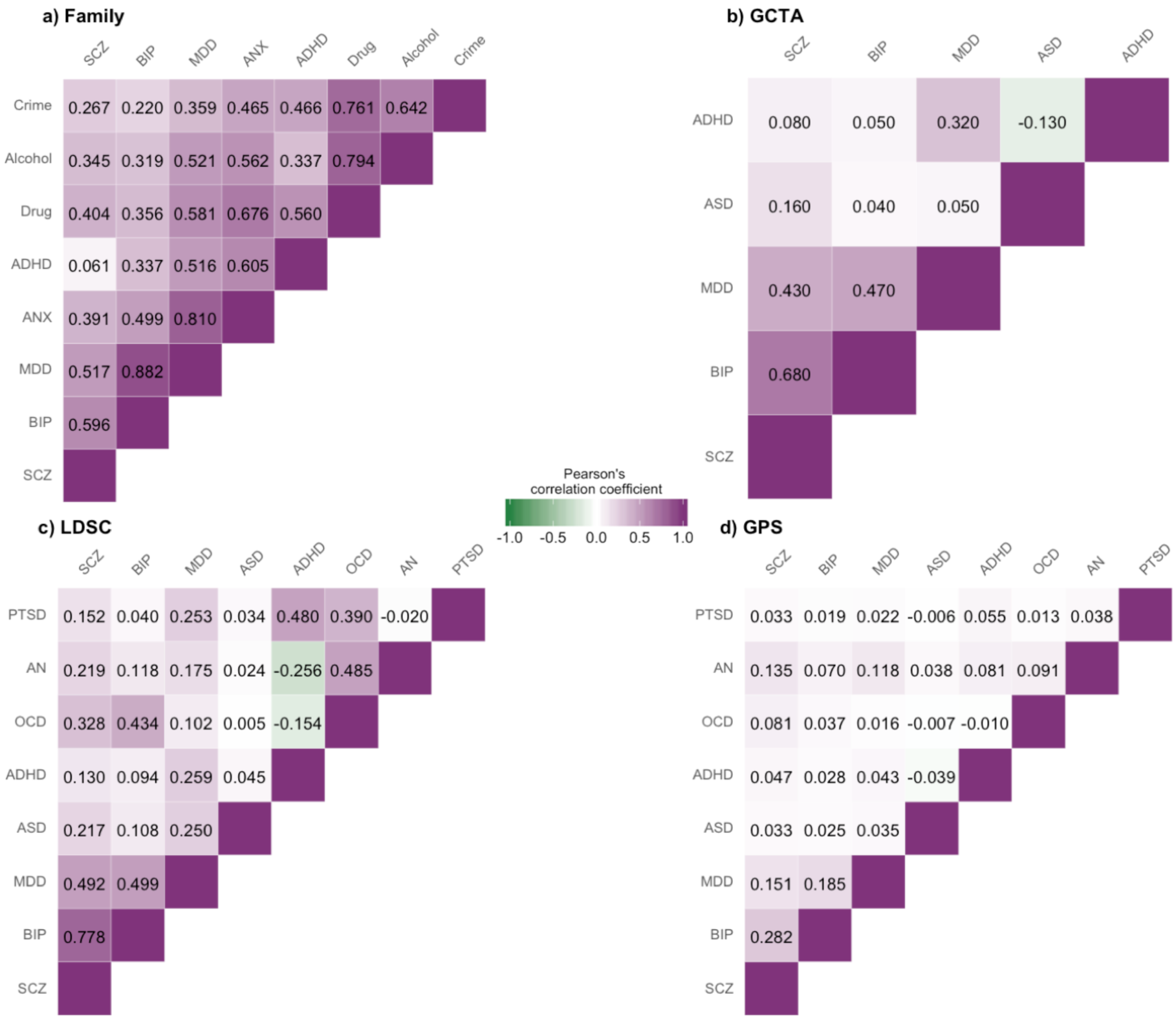
Genetic correlations from family analysis (a), Genome-wide Complex Trait Analysis (b), Linkage-Disequilibrium Score Regression (c) and Genome-wide Polygenic Score (d) analysis. Values represent Pearson’s correlation coefficients. SCZ = Schizophrenia; BIP = Bipolar Disorder; MDD = Major Depressive Disorder; ASD = Autism Spectrum Disorder; ADHD = Attention-Deficit/Hyperactivity Disorder; ANX = Anxiety; OCD = Obsessive-Compulsive Disorder; AN = Anorexia Nervosa; PTSD = Post-Traumatic Stress Disorder; Drug = Drug abuse; Alcohol = Alcohol abuse; Crime = Convictions of violent crimes.

### Principal Component Analysis

Principal component analyses provided converging evidence for a general psychopathology factor. Figure 2 shows that all four correlation matrices yielded a first unrotated principal component that accounted for a considerable amount of variance. The first principal component explained 57%, 43%, 34% and 19% in family, GCTA, LDSC and GPS data, respectively. (For proportion of variance explained by the other unrotated principal components, see Supplementary Table S4.)

**Figure 2.**
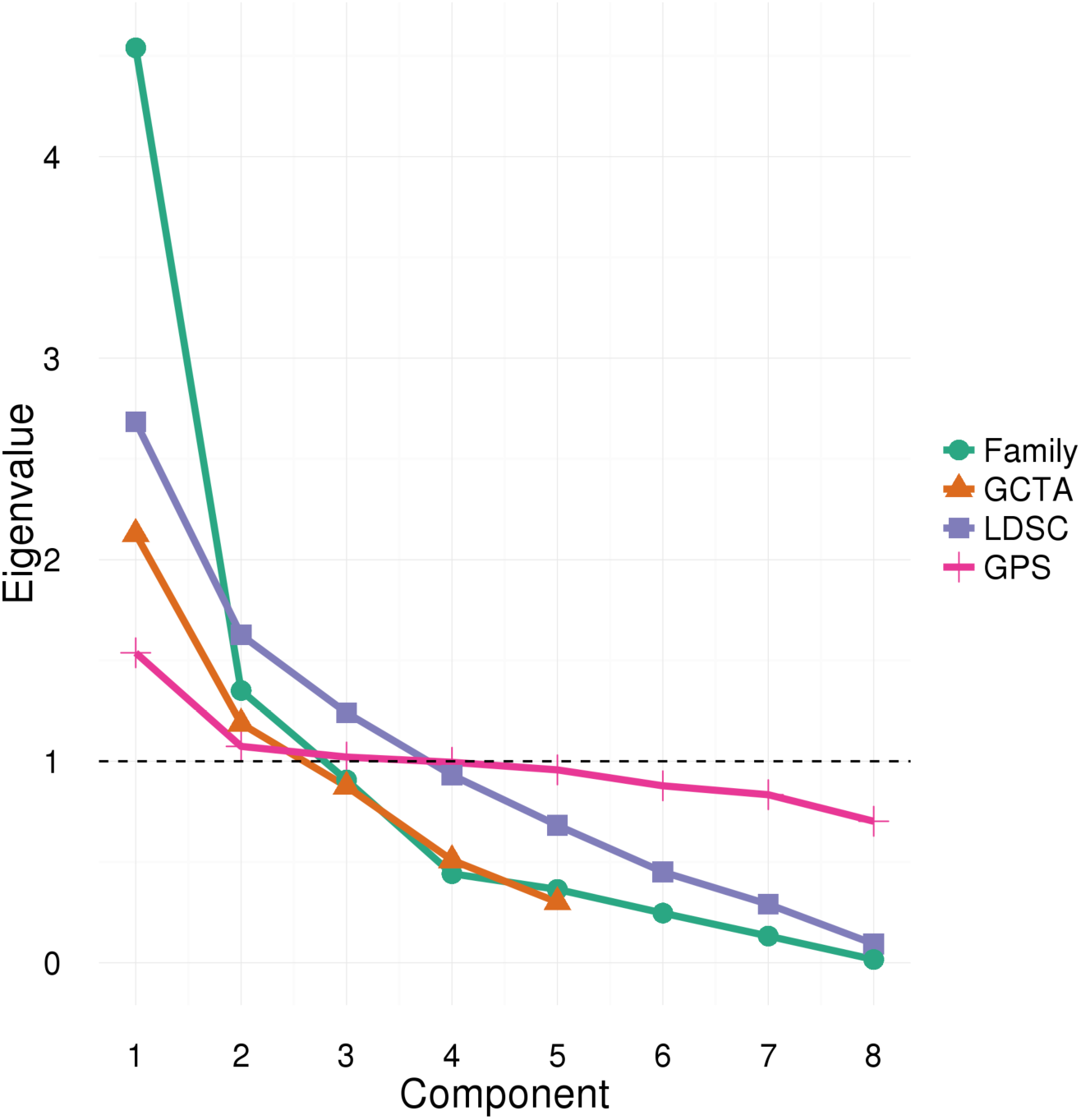
Scree plot showing eigenvalues for each principal component after performing PCA on genetic correlation matrices for four genetically sensitive methods: family analysis, Genome-wide Complex Trait Analysis (GCTA), Linkage-Disequilibrium Score Regression (LDSC) and Genome-wide Polygenic Scoring (GPS). The dashed line represents the cut-off for principal component retention based on the Kaiser’s λ > 1 criterion^28^.

Figure 3 shows first unrotated principal component loadings of all psychopathological traits for the four genetic methods. The loadings on the first unrotated principal component (PC) mirrored the genetic correlations (Figure 1): the average loadings were 0.75 for family data, 0.58 for GCTA, 0.53 for LDSC and 0.38 for GPS. We were able to test the statistical significance of loadings in family and GPS analyses, and found that all traits significantly loaded onto the first unrotated PC (all *p*-values ≤ 5.92×10^−24^), even though the GPS data showed some of the lowest loadings. The variation in factor loadings across the four methods can be explained by the inclusion of different disorders, as average loadings for the disorders in common were highly similar (family = 0.56; GCTA = 0.54; LDSC = 0.52; GPS = 0.42). Schizophrenia, Bipolar, and Depression consistently had the highest loadings on the first unrotated principal component across all genetic approaches. In contrast, Autistic Spectrum Disorder and Attention-Deficit/Hyperactivity Disorder were among the lowest-loading traits.

**Figure 3.**
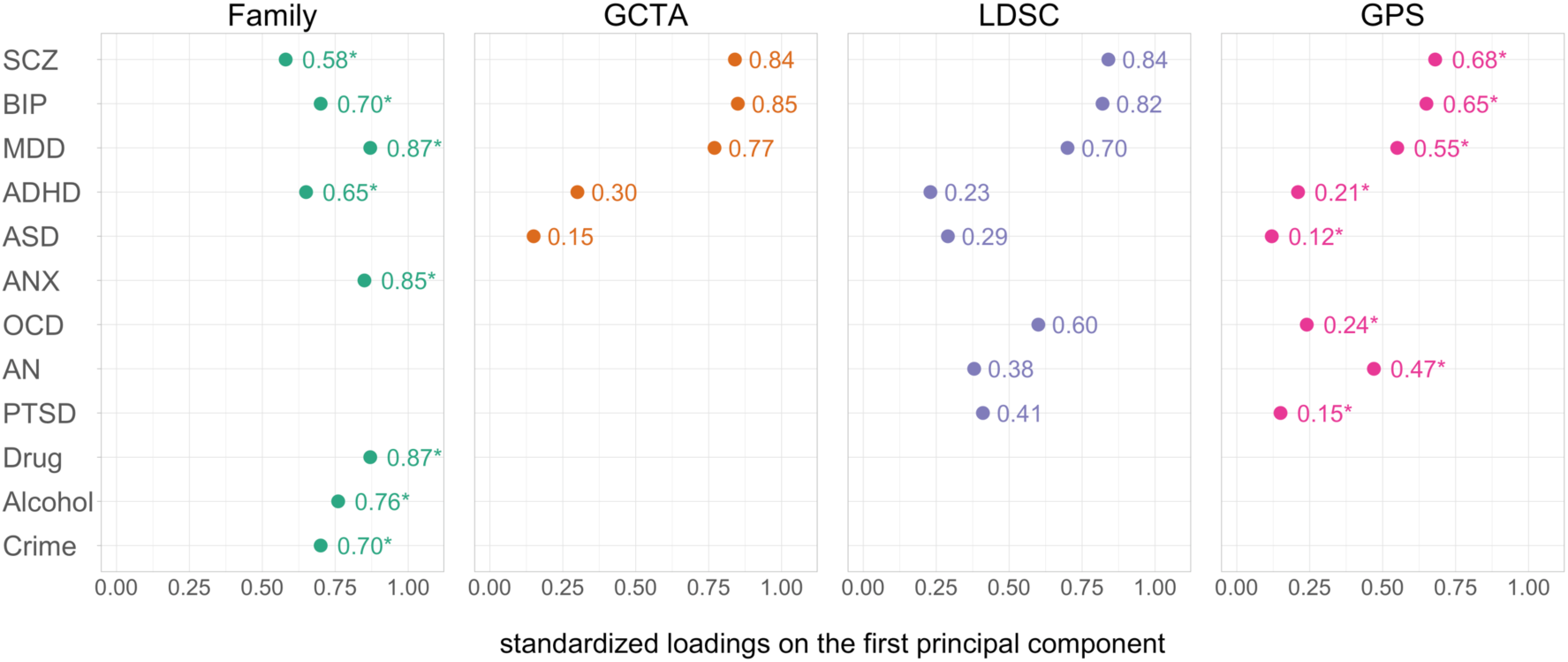
Loadings of psychopathology traits on the first unrotated principal component for each of the four types of genetic data. GCTA = Genome-wide Complex Trait Analysis; LDSC = Linkage-Disequilibrium Score Regression; GPS = Genome-wide Polygenic Score; SCZ = Schizophrenia; BIP = Bipolar Disorder; MDD = Major Depressive Disorder; ASD = Autism Spectrum Disorder; ADHD = Attention-Deficit/Hyperactivity Disorder; ANX = Anxiety; OCD = Obsessive-Compulsive Disorder; AN = Anorexia Nervosa; PTSD = Post-Traumatic Stress Disorder; Drug = Drug abuse; Alcohol = Alcohol abuse; Crime = Convictions of violent crimes. “*“ = reached statistical significance of *p* ≤ 5.92×10^−24^; it was only possible to test the statistical significance for the loadings relating to GPS and family data (see Methods section for details).

Based on the criteria described in the Methods section, we retained two principal components for rotation for family, GCTA and GPS data, and three principal components for LDSC data (for more details, see Supplementary Table S4). However, to improve comparability of the rotated factor solutions across the four genetic methods, we kept two PCs for the LDSC data. Results of the rotation of three components for LDSC data can be found in Supplementary Table S5.

Figure 4 lists the loadings for the first two rotated factors after performing oblique rotation. Rotated factor loadings for all methods (family, GCTA, LDSC, GPS) show that Schizophrenia, Bipolar and Depression consistently load highly onto the same factor. This is expected from the higher genetic intercorrelations between these traits for all methods (Figure 1). For the remaining psychiatric traits, results were less consistent when comparing family data to genomic data (GCTA, LDSC, GPS). In part, this reflects the traits included – most notably, a drug abuse/crime factor emerged from the family data because, unlike the other datasets, Drug Abuse, Alcohol Abuse and Violent Crime were included and created the first rotated factor. Anxiety also contributed to both the rotated factors. For the three genomic methods, the second factor primarily included Attention-Deficit/Hyperactivity Disorder and Post-Traumatic Stress Disorder.

**Figure 4.**
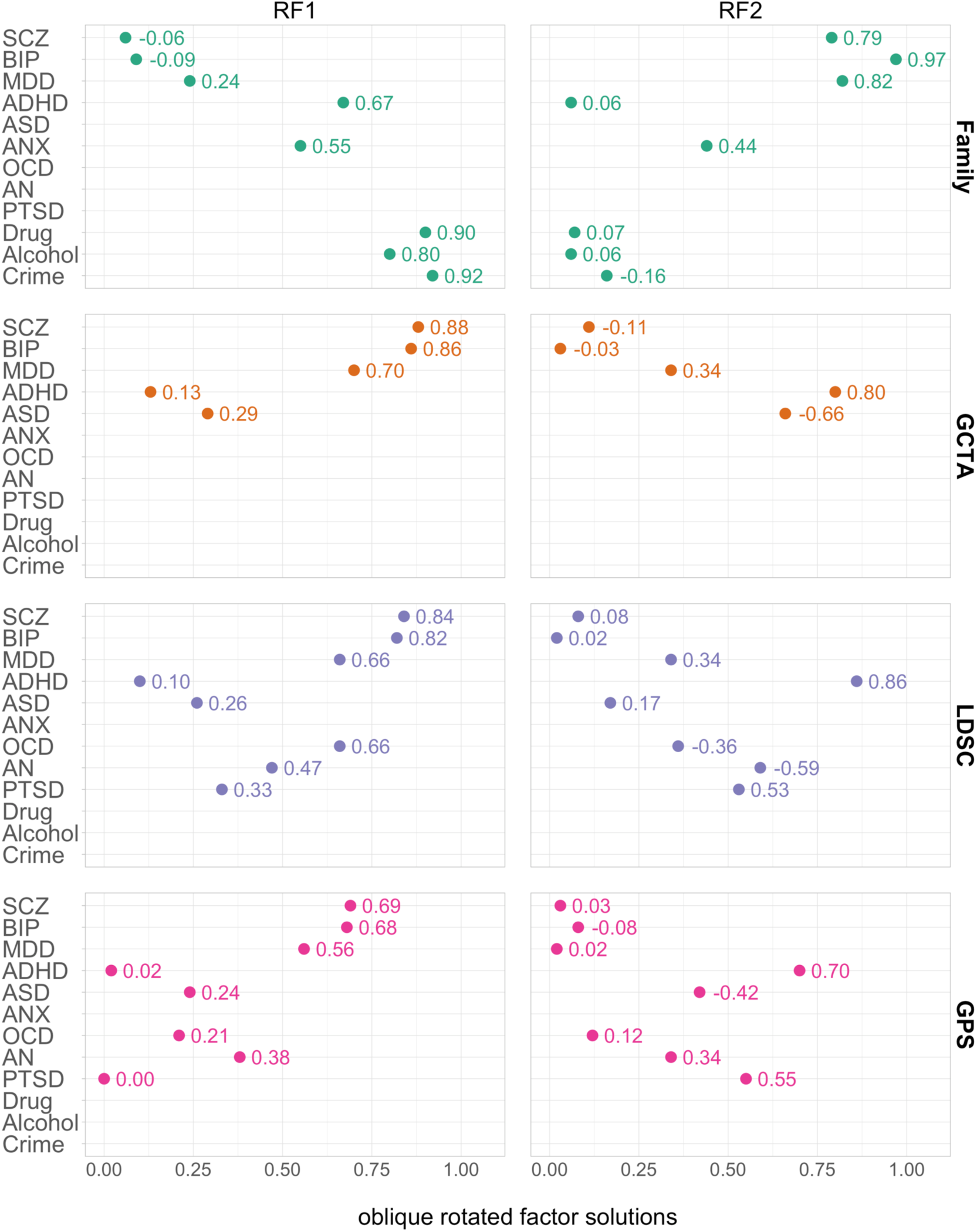
Rotated factor loadings for the four types of genetic data. RF = rotated factor based on oblique (*Oblimin*) rotation. GCTA = Genome-wide Complex Trait Analysis; LDSC = Linkage-Disequilibrium Score Regression; GPS = Genome-wide Polygenic Score; SCZ = Schizophrenia; BIP = Bipolar Disorder; MDD = Major Depressive Disorder; ASD = Autism Spectrum Disorder; ADHD = Attention-Deficit/Hyperactivity Disorder; ANX = Anxiety; OCD = Obsessive-Compulsive Disorder; AN = Anorexia Nervosa; PTSD = Post-Traumatic Stress Disorder; Drug = Drug abuse; Alcohol = Alcohol abuse; Crime = Convictions of violent crimes.

## Discussion

These results provide genetic support for *p*, a general factor of psychopathology that represents a single, continuous dimension of the psychiatric spectrum. The four methods used to estimate genetic correlations differ substantially: quantitative genetic analysis of siblings and half-siblings^15^, GCTA estimates based on SNP differences between unrelated individuals^17^, LDSC analysis based on GWA summary statistics, and GPS for individual data presented in this paper. Nonetheless, each of the principal component analyses from the four methods yielded a general factor on which all disorders loaded, explaining between 20% and 60% of the total variance.

Schizophrenia, Bipolar and Depression are the oldest and most consistently diagnosed psychiatric disorders, yet they are consistently among the highest-loading disorders on this genetic *p* factor. This finding is unlikely to be due to some artifact of genetic analysis because it was consistent across different genetic methods applied to different samples.

In contrast, Attention-Deficit/Hyperactivity Disorder and Autistic Spectrum Disorder consistently show the lowest loadings on the first unrotated principal component. An obvious hypothesis to account for this is that these two disorders are diagnosed in childhood, whereas the other disorders are diagnosed in adulthood. There is recent interest in diagnosing adult versions of these disorders^24,25^. If GWA studies were conducted on in adults, genetic correlations with other psychiatric disorders should be higher if this hypothesis is correct.

It is difficult to draw general conclusions about the other disorders that varied across the four genetic methods (Obsessive Compulsive Disorder, Anorexia, and Post-traumatic Stress Disorder, Anxiety, Drug Abuse, Alcohol Abuse, and Violent Crime). However, when any of these disorders were included in a study, they consistently contributed to a genetic *p* factor in the sense that they loaded on the first unrotated principal component.

Although the four genetic methods yielded similar patterns of correlations and patterns of loadings on the first unrotated principal component, they differed in the magnitude of their estimates of correlations and loadings, even when only considering the disorders in common (i.e. Schizophrenia, Bipolar, Depression, Autistic Spectrum Disorder). In principle, genetic correlations calculated through GCTA and LDSC should not differ substantially from family study estimates. Even though univariate SNP-h^2^ is generally lower than family-h^2^ because the estimate doesn’t include rare variants and nonadditive effects, this downward bias influences both nominator and denominator to equal extents when calculating genetic correlations (r_g_ = h_x_h_y_ / √h_x_^2^h_y_^2^), therefore cancelling out the bias^26^. However, if the correlation between causal SNPs is stronger for common variants than for rare variants, the SNP genetic correlation estimate would be higher than family study estimates, because only common SNPs are included in the analysis^19^. Nevertheless, for the disorders in common, family data produced higher average genetic correlations (0.49) than LDSC (0.34) and GCTA (0.38). An alternative explanation may be differing approaches to sample ascertainment and psychiatric diagnoses. In some genomic studies, sampling strategies may select ‘pure’ cases and exclude cases with other co-occuring conditions, and such ‘pure’ cases do not represent the disordered population^27^. In contrast, family data used in this study^15^ were based on a non-hierarchical approach to classification, thus allowing for greater overlap among the disorders.

GPS results yielded the lowest overall correlations, as it is arguably the most conceptually distinct method. A GPS is the aggregation of all genetic effects found in an independent GWA analysis in respect to an individual’s genotype. Therefore, GPS correlations index the extent to which the total variance of individuals’ GPS for one trait covaries with GPS for other traits. Two possible reasons why GPS correlations may be the lowest are that (i) in addition to true effects, a GPS includes the measurement error for all the SNPs tested across the genome in GWA analysis and (ii) a GPS is generated using genotypes from one cohort and effect sizes from a second, independent cohort.

What causes this genetic *p* factor? The positive manifold of the genetic *p* factor is agnostic about its causes. There are several, equally plausible hypotheses for the mechanisms that cause cross-disorder correlations^28^. One possible pathway may be *biological pleiotropy*, where DNA variants are causally involved in the development of several traits related to psychopathology. An alternative explanation is *mediated pleiotropy*, in which comorbidity occurs because DNA variants increase risk for one disorder, and then this disorder causes other disorders in turn. A third hypothesis is that DNA variants cause some general impairment that forms the core of various disorders, consequently producing genetic correlation between specific diagnoses. That is, the thousands of DNA variants associated with each symptom or disorder might affect all personality and cognitive processes that increase risk, thus providing many pathways to psychopathology.

Although it is remarkable how much genetic variance is explained by *p*, it does not explain all, or even most, of the genetic variance. Assuming a hierarchical model with *p* at the highest level^9,10^, broader psychiatric dimensions at a middle level, and specific psychopathologies at the lowest level, the question is how much genetic variance is accounted for by the three levels. In the realm of cognitive abilities, there continues to be debates about the nature of the middle level^29^. As compared to *p*, there is less clarity in our results about the nature of the second level of the hierarchical structure, as represented by the rotated factor solutions. One rotated factor consistently includes the psychotic disorders of Schizophrenia, Bipolar and Depression. However, the other rotated factor is less clear. For example, although Attention-Deficit/Hyperactivity Disorder loads on the second factor, it clusters positively with Post-Traumatic Stress Disorder in the LDSC and GPS results, positively with anxiety, substance abuse and crime in the family results, and negatively with Autistic Spectrum Disorder in the GCTA and GPS results. It may be that the second level of the hierarchical structure will remain unclear until analyses of this type begin to use a transdiagnostic approach, by beginning with symptoms and building a hierarchical model from the ground up. If these data become available in the future, we will be able test the genetic *p* factor model more formally by contrasting it to alternative models.

In addition to this limitation of our analyses, the primary limitation is ‘missing heritability’, the gap between SNP-h^2^ and family study heritability estimates. We used the most recent publicly available GWA summary statistics, some of which are considerably underpowered. This limitation most affects our GPS analyses, which predict genetic risk at the level of individuals. The modest SNP-h^2^ and measurement error of the GWA studies from which the GPS were derived are partly responsible for the low correlations between the GPS. More powerful GWA studies are in progress, and we are optimistic that new GPS will have improved predictive accuracy. More generally, GWA studies focused on phenotypic *p* should be able to capture genetic *p* to a greater extent than trying to derive genetic *p* from GWA studies of separate disorders that are sometimes diagnosed as ‘pure’ cases that exclude other diagnoses.

In conclusion, we report strong evidence for a genetic *p* factor that represents a continuous, underlying dimension of psychiatric risk using four distinct genetic methods. As GWA studies continue to increase in sample size as well as in the diversity of their target traits, our current results suggest that a genetic *p* factor could be useful in psychiatric research.

## Methods

### Sample

This study included 7,026 unrelated (i.e., one member per twin pair), genotyped individuals from the Twins Early Development Study (TEDS), a longitudinal birth cohort that recruited over 15,000 twin pairs between 1994-1996 who were born in England or Wales. Despite some attrition, the remaining cohort, as well as the genotyped subsample have been shown to represent the UK population^23,30^. Written informed consent was obtained from parents. Project approval was granted by King’s College London’s ethics committee for the Institute of Psychiatry, Psychology and Neuroscience (05.Q0706/228).

### Genome-wide Polygenic Scores (GPS)

To obtain individual-specific genetic measures for psychiatric traits, we created eight GPS in our independent sample of 7,026 individuals based on publicly available genome-wide association (GWA) summary statistics from the Psychiatric Genomics Consortium (PGC): Schizophrenia, Bipolar Disorder, Major Depressive Disorder, Autism Spectrum Disorder, Attention-Deficit/Hyperactivity Disorder, Obsessive-Compulsive Disorder, Anorexia Nervosa, Post-Traumatic Stress Disorder (Supplementary Table S1). Following quality control and imputation (see Supplementary Methods S1 for details), genotypic data included 515,100 genotyped or imputed SNPs (info=1). To calculate polygenic scores, we used a Bayesian approach, *LDpred*^31^, which modifies the summary statistic coefficients based on information on Linkage Disequilibrium (LD) and a prior on the effect size of each SNP. The final GPS is obtained as the sum of the trait-increasing alleles (each variant coded as 0,1, or 2), weighted by the posterior effect size estimates (for more information, see Supplementary Methods S2). All polygenic scores were adjusted for the first ten principal components of the genotype data, chip, batch and plate effects using the regression method. The resulting standardized residuals were used for subsequent analyses.

Based on polygenic scores for the eight psychopathology traits created in TEDS, we generated a correlation matrix for further use in the main analyses.

### Genetic correlations based on Linkage-Disequilibrium Score Regression (LDSC)

LDSC is a method used to estimate SNP-heritability (SNP-h^2^) based on GWA summary statistics only, and relies on the principle that the presence of LD in the study sample is correlated with the upward bias of GWA test statistics^18^. Cross-trait LDSC^19^ is an extension of this method and makes it possible to estimate the genetic relationship between two traits. For each SNP, this method establishes the covariance of the test statistics for trait *x* and trait *y*, and regresses this value on the LD score of that SNP (i.e. the sum of the squared correlations of the SNP with its surrounding SNPs), whereby the slope represents the genetic covariance. The genetic correlation is obtained by standardizing the covariance by the SNP-h^2^ for both traits (r_g_ = cov_xy_ / √h_x_^2^h_y_^2^).

We applied cross-trait LDSC analysis on the same eight PGC summary statistics used for polygenic score creation to generate a genetic correlation matrix for further analysis. (For univariate SNP-h^2^ results using LDSC, see Supplementary Table S2.)

### Genetic correlations based on Genome-wide Complex Trait Analysis (GCTA)

In addition to GPS and LDSC analysis, we also obtained genetic correlation matrices based on bivariate Genome-wide Complex Trait Analysis (GCTA)^32^. Unlike LDSC, which uses GWA summary statistics, GCTA requires individual-level genotype data and estimates genetic correlations from linear mixed model analysis by relating a pairwise genomic similarity matrix to a phenotypic covariance matrix between traits *x* and *y*. We used published GCTA genetic correlations^17^, which included five psychiatric disorders: Schizophrenia, Bipolar Disorder, Major Depressive Disorder, Autism Spectrum Disorder, and Attention-Deficit/Hyperactivity Disorder. (For univariate SNP-h^2^ estimates, see Supplementary Table S3.)

### Genetic correlations based on family data

Finally, we used genetic correlations based on quantitative genetic analysis comparing 3,475,122 Swedish full-siblings and half-siblings, who are genetically similar 50% and 25%, respectively, for additive genetic effects. The genetic correlations were not included in the original publication^15^ but were kindly prepared and shared by the lead author, Erik Pettersson of the Karolinska Institute. The analysis included seven psychopathology traits (Schizophrenia, Bipolar Disorder, Attention-Deficity/Hyperactivity Disoder, Major Depressive Disorder, Anxiety, Alcohol Use Disorder and Drug Abuse), as well as measured violent crimes convictions. Schizoaffective Disorder was redundant with Schizophrenia and thus omitted here (Supplementary Figure S1).

## Statistical Analyses

### Principal Component Analysis

To test the hypothesis that a general genetic *p* factor emerges from the genetic relationships among psychopathology traits, we performed eigenvalue decomposition through Principal Component Analysis (PCA), which aims to maximize variation of the first principal component (PC)^33^. We applied PCA to genetic correlation matrices derived from family analysis (8×8 matrix), GCTA (5×5 matrix), LDSC (8×8 matrix), and GPS analysis (8×8 matrix) to test whether a first principal component explains a substantial amount of the variance, and whether all psychiatric traits load onto this component.

We also tested the statistical significance of the factor loadings, which represent correlations between the original standardized variables and the factors. By calculating the t-statistic of the correlation coefficients, we were able to derive empirical *p*-values based on the *t*-statistic distribution with n−2 degrees of freedom^34^. Significance testing was applied to family and GPS loadings only because we were unable to obtain degrees of freedom for GCTA and LDSC data, which is required for the calculation of *t*.

The decision of how many components to retain for rotation was based on three criteria: (i) the Kaiser criterion^35^ of eigenvalue λ > 1; (ii) parallel analysis^36^, and (iii) scree plot inspection^37^ (for a more detailed description, see Supplementary Methods S3). To improve interpretability of the extracted components, we performed oblique rotation using the *Oblimin* method. We chose this approach, which permits factors to be correlated, because previous work using phenotypic data showed considerable associations between latent psychopathology dimensions^3,5^.

Analyses were performed in R^38^, using the *hornpa*^39^ package to perform parallel analysis, the *psych*^40^ package to conduct PCA (using the *principal* function), and the *GPArotation*^41^ package to apply oblique rotation.

## Acknowledgements

We gratefully acknowledge the ongoing contribution of the participants in the Twins Early Development Study (TEDS) and their families. The authors also wish to acknowledge Erik Pettersson for making available family study genetic correlation data for the use in this study. TEDS is supported by a program grant to RP from the UK Medical Research Council (MR/M021475/1 and previously G0901245), with additional support from the US National Institutes of Health (HD044454; HD059215). SS is supported by the MRC/IoPPN Excellence Award and by the EU Framework Programme 7 (602768). The research leading to these results has also received funding from the European Research Council under the European Union’s Seventh Framework Programme (FP7/2007-2013) / ERC grant agreement n° 295366. This study represents independent research part funded by the National Institute for Health Research (NIHR) Biomedical Research Centre at South London and Maudsley NHS Foundation Trust and King’s College London. The views expressed are those of the author(s) and not necessarily those of the NHS, the NIHR or the Department of Health. High performance computing facilities were funded with capital equipment grants from the GSTT Charity (TR130505) and Maudsley Charity (980).

## Author Contributions

Study concept and design: SS, RP. Processed and quality controlled genotype data: SS. Analysis of data: SS. Interpretation of data: All authors. Wrote the paper: SS, RP. Contributed to and critically reviewed the manuscript: All authors.

## References

1 Kessler RC, Chiu WT, Demler O, Walters EE. Prevalence, Severity, and Comorbidity of 12-Month DSM-IV Disorders in the National Comorbidity Survey Replication. Arch Gen Psychiatry 2005; 62: 617–627.

2 Kotov R, Krueger RF, Watson D, Achenbach TM, Althoff RR, Bagby RM et al. The Hierarchical Taxonomy of Psychopathology (HiTOP): A dimensional alternative to traditional nosologies. Journal of Abnormal Psychology 2017; 126: 454–477.

3 Wright AGC, Krueger RF, Hobbs MJ, Markon KE, Eaton NR, Slade T. The structure of psychopathology: Toward an expanded quantitative empirical model. Journal of Abnormal Psychology 2013; 122: 281–294.

4 Lahey BB, Applegate B, Hakes JK, Zald DH, Hariri AR, Rathouz PJ. Is there a general factor of prevalent psychopathology during adulthood? Journal of Abnormal Psychology 2012; 121: 971–977.

5 Caspi A, Houts RM, Belsky DW, Goldman-Mellor SJ, Harrington H, Israel S et al. The p Factor. Clinical Psychological Science 2014; 2: 119–137.

6 Spearman C. ‘General intelligence’ objectively determined and measured. American Journal of Psychology 1904; 15: 201–292.

7 Carroll JB. Human Cognitive Abilities: A Survey of Factor-Analytic Studies. Cambridge University Press: Cambridge, 1993

8 Deary IJ. Intelligence. Annual Review of Psychology 2012; 63: 453–482.

9 Caspi A, Moffitt TE. All for one and one for all: Mental disorders in one dimension. American Journal of Psychiatry in press.

10 Lahey BB, Krueger RF, Rathouz PJ, Waldman ID, Zald DH. A Hierarchical Causal Taxonomy of Psychopathology Across the Life Span. Psychol Bull 2017; 143: 142–186.

11 McLaughlin KA, Gadermann AM, Hwang I, Sampson NA, Al-Hamzawi A, Andrade LH et al. Parent psychopathology and offspring mental disorders: Results from the WHO World Mental Health Surveys. British Journal of Psychiatry 2012; 200: 290–299.

12 Kendler KS. Major depression and generalised anxiety disorder - Same genes, (partly) different environments - Revisited. British Journal of Psychiatry 1996; 168: 68–75.

13 Lichtenstein P, Yip BH, Björk C, Pawitan Y, Cannon TD, Sullivan PF et al. Common genetic determinants of schizophrenia and bipolar disorder in Swedish families: a population-based study. The Lancet 2009; 373: 234–239.

14 Kendler KS, Aggen SH, Knudsen GP, Røysamb E, Neale MC, Reichborn-Kjennerud T. The Structure of Genetic and Environmental Risk Factors for Syndromal and Subsyndromal Common DSM-IV Axis I and All Axis II Disorders. American Journal of Psychiatry 2011; 168: 29–39.

15 Pettersson E, Larsson H, Lichtenstein P. Common psychiatric disorders share the same genetic origin: a multivariate sibling study of the Swedish population. Molecular Psychiatry 2015 21:5 2016; 21: 717–721.

16 Cross-Disorder Group of the Psychiatric Genomics Consortium. Identification of risk loci with shared effects on five major psychiatric disorders: a genome-wide analysis. The Lancet 2013; 381: 1371–1379.

17 Lee SH, Ripke S, Neale BM, Faraone SV, Purcell SM, Perlis RH et al. Genetic relationship between five psychiatric disorders estimated from genome-wide SNPs. Nat Genet 2013; 45: 984–994.

18 Bulik-Sullivan BK, Loh P-R, Finucane HK, Ripke S, Yang J, Patterson N et al. LD Score regression distinguishes confounding from polygenicity in genome-wide association studies. Nat Genet 2015; 47: 291–295.

19 Bulik-Sullivan B, Finucane HK, Anttila V, Gusev A, Day FR, Loh P-R et al. An atlas of genetic correlations across human diseases and traits. Nat Genet 2015; 47: 1236–1241.

20 Anttila V, Bulik-Sullivan B, Finucane HK, Bras J, Duncan L, Escott-Price V, et al. Analysis of shared heritability in common disorders of the brain. bioRxiv 2016; : 048991.

21 Krapohl E, Euesden J, Zabaneh D, Pingault J-B, Rimfeld K, Stumm von S et al. Phenome-wide analysis of genome-wide polygenic scores. Molecular Psychiatry 2015 21:5 2016; 21: 1188–1193.

22 Dudbridge F. Polygenic Epidemiology. Genetic Epidemiology 2016; 40: 268–272.

23 Haworth CMA, Davis OSP, Plomin R. Twins Early Development Study (TEDS): A Genetically Sensitive Investigation of Cognitive and Behavioral Development From Childhood to Young Adulthood. Twin Res Hum Genet 2013; 16: 117–125.

24 Asherson P, Buitelaar J, Faraone SV, Rohde LA. Adult attention-deficit hyperactivity disorder: key conceptual issues. The Lancet Psychiatry 2016; 3: 568–578.

25 Poon KK, Sidhu DJK. Adults with autism spectrum disorders: a review of outcomes, social attainment, and interventions. Current Opinion in Psychiatry 2017; 30: 77–84.

26 Trzaskowski M, Davis OSP, DeFries JC, Yang J, Visscher PM, Plomin R. DNA Evidence for Strong Genome-Wide Pleiotropy of Cognitive and Learning Abilities. Behav Genet 2013; 43: 267–273.

27 Newman DL, Moffitt TE, Caspi A, Silva PA. Comorbid mental disorders: implications for treatment and sample selection. Journal of Abnormal Psychology 1998; 107: 305–311.

28 Solovieff N, Cotsapas C, Lee PH, Purcell SM, Smoller JW. Pleiotropy in complex traits: challenges and strategies. Nat Rev Genet 2013; 14: 483–495.

29 Johnson W, Bouchard T. The structure of human intelligence: It is verbal, perceptual, and image rotation (VPR), not fluid and crystallized. Intelligence 2005; 33: 393–416.

30 Selzam S, Krapohl E, Stumm von S, O’Reilly PF, Rimfeld K, Kovas Y et al. Predicting educational achievement from DNA. Molecular Psychiatry 2015 21:5 2017; 22: 267–272.

31 Vilhjalmsson BJ, Yang J, Finucane HK, Gusev A, Lindstrom S, Ripke S et al. Modeling Linkage Disequilibrium Increases Accuracy of Polygenic Risk Scores. Am J Hum Genet 2015; 97: 576–592.

32 Lee SH, Yang J, Goddard ME, Visscher PM, Wray NR. Estimation of pleiotropy between complex diseases using single-nucleotide polymorphism-derived genomic relationships and restricted maximum likelihood. Bioinformatics 2012; 28: 2540–2542.

33 Jolliffe IT. Principal Component Analysis and Factor Analysis. In: Principal Component Analysis. Springer: New York, NY, 1986, pp 115–128.

34 Yamamoto H, Fujimori T, Sato H, Ishikawa G, Kami K, Ohashi Y. Statistical hypothesis testing of factor loading in principal component analysis and its application to metabolite set enrichment analysis. BMC Bioinformatics 2014; 15. doi:10.1186/1471-2105-15-51.

35 Kaiser HF. The Application of Electronic Computers to Factor Analysis. Educational and Psychological Measurement 1960; 20: 141–151.

36 Horn JL. A rationale and test for the number of factors in factor analysis. Psychometrika 1965; 30: 179–185.

37 Cattell RB. The Scree Test For The Number Of Factors. Multivariate Behav Res 1966; 1: 245–276.

38 R Core Team. R: A Language and Environment for Statistical Computing. R Foundation for Statistical Computing 2017. https://www.r-project.org.

39 Huang F. hornpa: Horn’s (1965) Test to Determine the Number of Components/Factors 2015. https://CRAN.R-project.org/package=hornpa.

40 Revelle WR. psych: Procedures for Personality and Psychological Research. 2017.https://CRAN.R-project.org/package=psych.

41 Bernaards CA, Jennrich RI. Gradient Projection Algorithms and Software for Arbitrary Rotation Criteria in Factor Analysis. Educational and Psychological Measurement 2016; 65: 676–696.

