## Supplementary Materials for "A polygenic *p* factor for major psychiatric disorders"

#### Title

### Supplementary Tables

#### Table S1. Publicly available PGC GWA studies used for creation of polygenic scores

| Trait | Cases | Controls | Total Sample | SNPs in common[a] | Year Published | PMID | Reference |
|---|---|---|---|---|---|---|---|
| SCZ | 36989 | 113075 | 150064 | 499197 | 2014 | 25056061 | 1 |
| BIP | 7481 | 9250 | 16731 | 409796 | 2011 | 21926972 | 2 |
| MDD | 9240 | 9519 | 18759 | 374208 | 2013 | 22472876 | 3 |
| ASD | 7387 | 8567 | 15954 | 456246 | 2017 | 28540026 | 4 |
| ADHD | 20183 | 35191 | 55374 | 469352 | 2017 | NA | 5 |
| OCD | 2688 | 7037 | 9725 | 498602 | 2017 | 28761083 | 6 |
| AN | 3495 | 10982 | 14477 | 477354 | 2017 | 28494655 | 7 |
| PTSD | 2424 | 7113 | 9537 | 499383 | 2017 | 28439101 | 8 |

Note. [a]= number of SNPs in common between GWA summary statistics and individual-level genotypes in target sample. SCZ = Schizophrenia, BIP = Bipolar Disorder, MDD = Major Depressive Disorder, ASD = Autism Spectrum Disorder, ADHD = Attention-Deficit/Hyperactivity Disorder, OCD = Obsessive-Compulsive Disorder, AN = Anorexia Nervosa, PTSD = Post-Traumatic Stress Disorder.

Table S2. Liability scale univariate SNP-heritability estimates based on LD-Score Regression

| **Trait** | **$h^2$** | **se** | ***z*-$h^2$** | **Lambda GC** | **Mean $Chi^2$** | **Intercept** | **Population prev** | **Sample prev** |
|---|---|---|---|---|---|---|---|---|
| SCZ | 0.179 | 0.007 | 24.792 | 1.592 | 1.783 | 1.038 | 0.010 | 0.250 |
| BIP | 0.250 | 0.023 | 10.810 | 1.159 | 1.183 | 1.015 | 0.010 | 0.450 |
| MDD | 0.188 | 0.036 | 5.208 | 1.074 | 1.077 | 1.013 | 0.150 | 0.490 |
| ASD | 0.186 | 0.025 | 7.504 | 1.071 | 1.081 | 0.987 | 0.010 | 0.460 |
| ADHD | 0.211 | 0.014 | 15.409 | 1.250 | 1.297 | 1.033 | 0.050 | 0.360 |
| OCD | 0.243 | 0.038 | 6.460 | 1.053 | 1.056 | 0.993 | 0.015 | 0.280 |
| AN | 0.182 | 0.029 | 6.313 | 1.077 | 1.079 | 1.009 | 0.010 | 0.240 |
| PTSD | 0.132 | 0.059 | 2.253 | 1.017 | 1.013 | 0.994 | 0.080 | 0.250 |

Note. $h^2$ = SNP-heritability derived from LD-Score Regression. Z- $h^2$ = z-score heritability ($h^2$/se). Intercept = LDSC intercept. prev = prevalences used to calculate $h^2$. Sample prevalence = cases/(cases+controls) based on GWAS. SCZ = Schizophrenia, BIP = Bipolar Disorder, MDD = Major Depressive Disorder, ASD = Autism Spectrum Disorder, ADHD = Attention-Deficit/Hyperactivity Disorder, OCD = Obsessive-Compulsive Disorder, AN = Anorexia Nervosa, PTSD = Post-Traumatic Stress Disorder.

Table S3. Liability scale univariate SNP-heritability estimates based on Genome-wide Complex Trait Analysis

| **Trait** | **$h^2$** | **se** | **z-$h^2$** | **Cases** | **Controls** | **Population prev** | **Sample prev** |
|---|---|---|---|---|---|---|---|
| SCZ | 0.230 | 0.008 | 28.750 | 9087 | 12171 | 0.01 | 0.43 |
| BIP | 0.250 | 0.012 | 20.833 | 6704 | 9031 | 0.01 | 0.43 |
| MDD | 0.210 | 0.021 | 10.000 | 9041 | 9381 | 0.15 | 0.49 |
| ASD | 0.170 | 0.025 | 6.800 | 3303 | 3428 | 0.01 | 0.46 |
| ADHD | 0.280 | 0.023 | 12.174 | 4163 | 12040 | 0.05 | 0.26 |

Note. $h^2$ = SNP-heritability derived from GCTA as reported in[9] . Z-$h^2$ = z-score heritability ($h^2$/se). prev = prevalences used to calculate $h^2$. Sample prevalence = cases/(cases+controls) based on study sample. SCZ  = Schizophrenia, BIP = Bipolar Disorder, MDD = Major Depressive Disorder, ASD = Autism Spectrum Disorder, ADHD = Attention-Deficit/Hyperactivity Disorder.

Table S4. Results from parallel analysis and initial solutions from unrotated Principal Component Analysis

| | | PC1 | PC2 | PC3 | PC4 | PC5 | PC6 | PC7 | PC8 |
|---|---|---|---|---|---|---|---|---|---|
| **Family analysis** | Parallel Analysis λ[a] | 1.003 | 1.002 | 1.001 | 1.001 | 1.000 | 1.000 | 0.999 | 0.999 |
| | λ | **4.54** | **1.35** | 0.91 | 0.44 | 0.36 | 0.25 | 0.13 | 0.02 |
| | Proportion of Variance | **0.57** | **0.17** | 0.11 | 0.06 | 0.05 | 0.03 | 0.02 | 0.00 |
| | Cumulative Variance | **0.57** | **0.74** | 0.85 | 0.90 | 0.95 | 0.98 | 1.00 | 1.00 |
| **GCTA** | Parallel Analysis λ | - | - | - | - | - | - | - | - |
| | λ | **2.13** | **1.19** | 0.87 | 0.51 | 0.3 | - | - | - |
| | Proportion of Variance | **0.43** | **0.24** | 0.17 | 0.1 | 0.06 | - | - | - |
| | Cumulative Proportion | **0.43** | **0.66** | 0.84 | 0.94 | 1.00 | - | - | - |
| **LDSC** | Parallel Analysis λ | - | - | - | - | - | - | - | - |
| | λ | **2.68** | **1.63** | **1.24** | 0.93 | 0.68 | 0.45 | 0.29 | 0.09 |
| | Proportion of Variance | **0.34** | **0.2** | **0.16** | 0.12 | 0.09 | 0.06 | 0.04 | 0.01 |
| | Cumulative Proportion | **0.34** | **0.54** | **0.69** | 0.81 | 0.90 | 0.95 | 0.99 | 1.00 |
| **GPS** | Parallel Analysis λ[b] | **1.065** | **1.044** | 1.028 | 1.014 | 1.003 | 0.991 | 0.978 | 0.964 |
| | λ | **1.54** | **1.07** | 1.02 | 0.99 | 0.96 | 0.88 | 0.83 | 0.70 |
| | Proportion of Variance | **0.19** | **0.13** | 0.13 | 0.12 | 0.12 | 0.11 | 0.10 | 0.09 |
| | Cumulative Proportion | **0.19** | **0.33** | 0.45 | 0.58 | 0.7 | 0.81 | 0.91 | 1.00 |

Note. λ = Eigenvalue. [a] Parameters used for parallel analysis: sample size = 3,475,112, variables = 8, repetitions = 1,000. [b] Parameters used for parallel analysis: sample size = 7,026, variables = 8, repetitions = 1,000. PCs in bold represent the PCs that passed the parallel analysis selection criteria[9].

Table S5. Rotated factor loadings based on three factors for LDSC data

| | RF1 | RF2 | RF3 |
|---|---|---|---|
| SCZ | 0.86 | 0.28 | -0.02 |
| BIP | 0.84 | 0.14 | -0.06 |
| MDD | 0.75 | -0.11 | 0.17 |
| ADHD | 0.15 | -0.39 | 0.77 |
| ASD | 0.52 | -0.30 | -0.15 |
| OCD | 0.12 | 0.86 | 0.19 |
| AN | 0.10 | 0.74 | -0.21 |
| PTSD | -0.05 | 0.25 | 0.91 |
| Factor | 1.00 | | |
| Correlations | 0.20 | 1.00 | |
| | 0.18 | 0.00 | 1.00 |

Note. RF = rotated factor based on oblique (*Oblimin*) rotation. SCZ = Schizophrenia, BIP = Bipolar Disorder, MDD = Major Depressive Disorder, ASD = Autism Spectrum Disorder, ADHD = Attention-Deficit/Hyperactivity Disorder, OCD = Obsessive-Compulsive Disorder, AN = Anorexia Nervosa, PTSD = Post-Traumatic Stress Disorder.

### Supplementary Figures

Figure S1. Original genetic correlation matrix including schizoaffective disorder as derived from family analysis

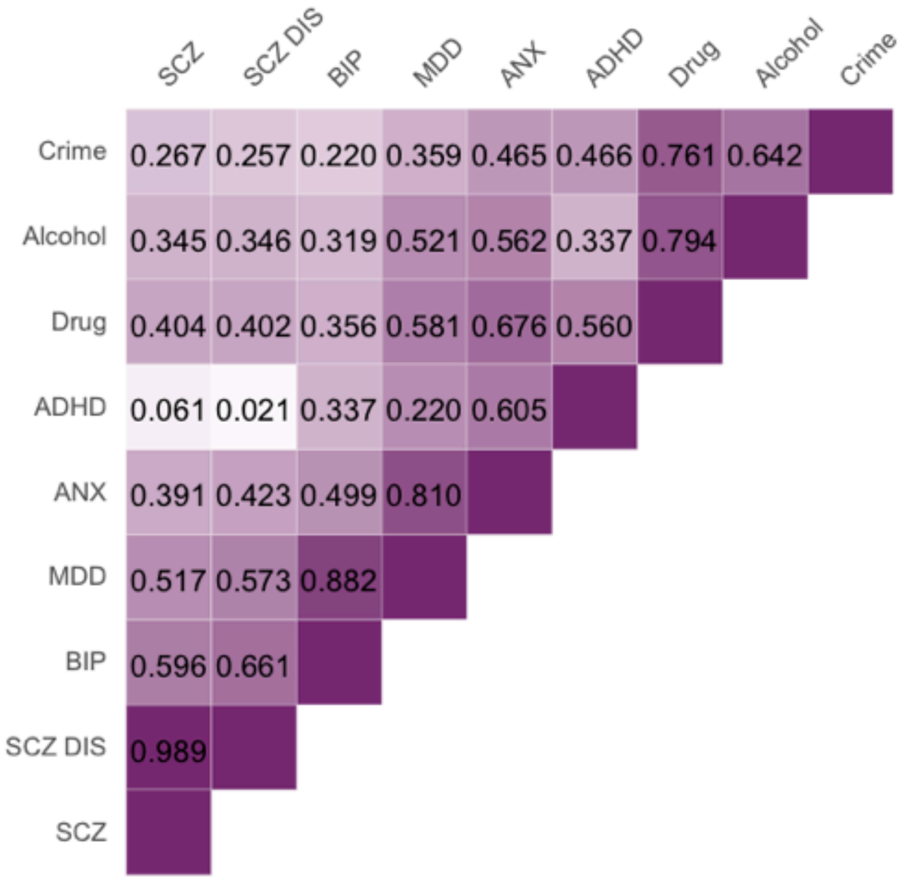

Note. SCZ = Schizophrenia, SCZ DIS = Schizoaffective Disorder, BIP = Bipolar Disorder, MDD = Major Depressive Disorder, ANX = Anxiety, ADHD = Attention-Deficit/Hyperactivity Disorder, Drug = Drug abuse, Alcohol = Alcohol abuse, Crime = Convictions of Violent Crimes

### Supplementary Methods

#### Methods S1. Genotyping and Quality Control

DNA for 8,122 individuals was extracted from saliva and buccal cheek swab samples and hybridized to HumanOmniExpressExome-8v1.2 genotyping arrays at the Institute of Psychiatry, Psychology and Neuroscience Genomics & Biomarker Core Facility. The raw image data from the array were normalized, pre-processed, and filtered in GenomeStudio according to Illumina Exome Chip SOP v1.4. (http://confluence.brc.iop.kcl.ac.uk:8090/display/PUB/Production+Version%3A+Illumina+Exome+Chip+SOP+v1.4). In addition, prior to genotype calling, 919 multi-mapping SNPs and 501 samples with call rate <0.95 were removed. The ZCALL program (see Web resources section) was used to augment the genotype calling for samples and SNPs that passed the initial QC.

DNA from 3,747 samples was extracted from buccal cheek swabs and genotyped at Affymetrix, Santa Clara, California, USA. From this sample, 3,665 samples were successfully hybridized to AffymetrixGeneChip 6.0 SNP genotyping arrays (http://www.affymetrix.com/support/technical/datasheets/genomewide_snp6_datasheet.pdf) using experimental protocols recommended by the manufacturer (Affymetrix Inc., Santa Clara, CA). The raw image data from the arrays were normalized and pre-processed at the Wellcome Trust Sanger Institute, Hinxton, UK for genotyping as part of the Wellcome Trust Case Control Consortium 2 (https://www.wtccc.org.uk/ccc2/) according to the manufacturer's guidelines (http://www.affymetrix.com/support/downloads/manuals/genomewidesnp6_manual.pdf). Genotypes for the Affymetrix arrays were called using CHIAMO (https://mathgen.stats.ox.ac.uk/genetics_software/chiamo/chiamo.html).

After initial quality control and genotype calling, the same quality control was performed on the samples genotyped on the Illumina and Affymetrix platforms separately using PLINK[10,11], R[12], BCFtools[13] and EIGENSOFT[14,15].

Samples were removed from subsequent analyses on the basis of call rate (<0.98), suspected non-European ancestry, heterozygosity, and relatedness other than dizygotic twin status. SNPs were excluded if the minor allele frequency was smaller than 0.5%, if more than 2% of genotype data were missing, or if the Hardy Weinberg *p*-value was lower than $10^{-5}$. Non-autosomal markers and indels were removed. Association between SNP and the platform, batch, plate or well on which samples were genotyped was calculated; SNPs with an effect *p*-value < $10^{-4}$ were excluded. The final sample comprised of 10,346 samples, including 7,026 unrelated individuals from which 3,320 individuals had a genotyped dizygotic co-twin. After quality control, genotype data included 4,776 individuals and 559,772 SNPs for the Illumina array, and 2,250 individuals and 635,269 SNPs for the Affymetrix array.

Genotypes from the two platforms were separately phased using EAGLE2[16], and imputed into the Haplotype Reference Consortium (release 1.1) using the Positional Burrows-Wheeler Transform method[17] through the Sanger Imputation Service. Prior to merging, we excluded variants with info <0.75 and removed non-overlapping SNPs between platforms. After merging, we tested for minor allele frequency differences between platforms and removed SNPs with an effect p-value < $10^{-4}$, and Hardy Weinberg p-value > $10^{-5}$. Using these criteria, 7,363,646 genotyped and well-imputed SNPs were retained for the analyses. In the present study, we included unrelated individuals only (N = 7,026). To ease high computational demands by the software LDpred[18] for polygenic scoring in large samples, we further excluded SNPs with info <1, leaving 515,100 SNPs for analysis.

We performed principal component analysis on a subset of 39,353 common (MAF > 5%), perfectly imputed (info = 1) autosomal SNPs, after stringent pruning to remove markers in linkage disequilibrium ($r^2 > 0.1$) and excluding high linkage disequilibrium genomic regions so as to ensure that only genome-wide effects were detected.

Methods S2. Polygenic score creation using LDpred

To calculate polygenic scores, we used a Bayesian approach, *LDpred* (version 0.9.0; https://github.com/bvilhjal/ldpred/blob/master/ldpred/LDpred.py)[18]. This method has been shown to outperform predictive accuracy of the conventional clumping and *p*-value thresholding approach[18]. Using LDpred, a posterior effect size for each SNP is derived by re-weighting the original summary statistic coefficient based on (i) the relative influence of a SNP given its level of LD with surrounding SNPs, and (ii) a prior on the effect size of each SNP. This prior is dependent on the heritability of the trait, as well as the fraction of markers assumed to causally influence the trait. The final GPS is obtained as the sum of the trait-increasing alleles (each variant coded as 0,1, or 2), weighted by the posterior effect size estimates. In contrast to clumping and thresholding, LDpred retains all the SNPs in the polygenic score that are common between GWA summary statistics and genotype data in the target sample.

LDpred is computationally demanding, especially in large sample sizes with a large number of SNPs. Therefore, we restricted our analyses to 515,100 SNPs that were perfectly imputed (info score of 1). Because levels of LD are considerably higher in relatives than in unrelated individuals[19], we only used genotypes of unrelated individuals to estimate LD structure in our sample. We applied a causal fraction of 1, which assumes that all SNPs contribute to the development of the trait. We decided on using this parameter to improve comparability with the other genetic methods used in this study (family analysis, GCTA, LDSC), which do not apply assumptions on the number of causally influencing SNPs and consider all genetic variations together in their estimation of genetic correlations.

#### Methods S3. Component selection criteria

According to the widely used Kaiser criterion[20], each PC with an eigenvalue $\lambda > 1$ represents an axis that explains more variance than a single variable itself, suggesting the retention of the component. However, randomly, uncorrelated data can produce components with $\lambda > 1$ due to chance covariation[21]. Therefore, we used parallel analysis[9] as the main criterion where possible. This method relies on the random generation of independent data, based on the same parameters as the original data (sample size; number of variables). To pass the parallel analysis criterion, eigenvalues from the study data must be larger than the 95$^{th}$ percentile of the distribution of the simulated components[22]. We performed PCAs to decompose four correlation matrices, one for each genetic method. Because LDSC and GCTA use thousands of SNPs to generate genetic correlations, we could only apply parallel analysis to family and GPS data, where the *n* number of variables used to generate a correlation matrix equates the number of variables obtained in the $n \times n$ correlation matrix. For analysis of GCTA and LDSC data, we therefore used the $\lambda > 1$ criterion instead. In addition, we performed scree plot inspection[23] to identify the point of inflection where the gradient of the line changes to a levelling-off slope, which signals that the components represent random variation rather than meaningful information.

### References

1 Schizophrenia Working Group of the Psychiatric Genomics Consortium. Biological insights from 108 schizophrenia-associated genetic loci. *Nature* 2014; **511**: 421–427.

2 Psychiatric GWAS Consortium Bipolar Disorder Working Group. Large-scale genome-wide association analysis of bipolar disorder identifies a new susceptibility locus near ODZ4. *Nat Genet* 2011; **43**: 977–983.

3 Major Depressive Disorder Working Group Consortium, Ripke S, Wray NR, Lewis CM, Hamilton SP, Weissman MM *et al.* A mega-analysis of genome-wide association studies for major depressive disorder. *Molecular Psychiatry 2015 21:5* 2013; **18**: 497–511.

4 The Autism Spectrum Disorders Working Group of The Psychiatric Genomics Consortium. Meta-analysis of GWAS of over 16,000 individuals with autism spectrum disorder highlights a novel locus at 10q24.32 and a significant overlap with schizophrenia. *Molecular Autism* 2017; **8**: 21.

5 Demontis D, Walters RK, Martin J, Mattheisen M, Als TD, Agerbo E *et al.* Discovery Of The First Genome-Wide Significant Risk Loci For ADHD. *bioRxiv* 2017; : 145581.

6 Arnold PD, Askland KD, Barlassina C, Bellodi L, Bienvenu OJ, Black D *et al.* Revealing the complex genetic architecture of obsessive–compulsive disorder using meta-analysis. *Molecular Psychiatry 2015 21:5* 2017; **1**: 457.

7 Duncan L, Yilmaz Z, Gaspar H, Walters R, Goldstein J, Anttila V *et al.* Significant Locus and Metabolic Genetic Correlations Revealed in Genome-Wide Association Study of Anorexia Nervosa. *American Journal of Psychiatry* 2017; **174**: 850–858.

8 Duncan LE, Ratanatharathorn A, Aiello AE, Almli LM, Amstadter AB, Ashley-Koch AE *et al.* Largest GWAS of PTSD (N=20 070) yields genetic overlap with schizophrenia and sex differences in heritability. *Molecular Psychiatry 2015 21:5* 2018; **23**: 666.

9 Horn JL. A rationale and test for the number of factors in factor analysis. *Psychometrika* 1965; **30**: 179–185.

10 Chang CC, Chow CC, Tellier LC, Vattikuti S, Purcell SM, Lee JJ. Second-generation PLINK: rising to the challenge of larger and richer datasets. *GigaScience 2015 4:1* 2015; **4**: 7.

11 Purcell S, Neale B, Todd-Brown K, Thomas L, Ferreira MAR, Bender D *et al.* PLINK: A Tool Set for Whole-Genome Association and Population-Based Linkage Analyses. *Am J Hum Genet* 2007; **81**: 559–575.

12 R Core Team. R: A Language and Environment for Statistical Computing. R Foundation for Statistical Computing 2017. https://www.r-project.org.

13 Li H. A statistical framework for SNP calling, mutation discovery, association mapping and population genetical parameter estimation from sequencing data. *Bioinformatics* 2011; **27**: 2987–2993.

14 Patterson N, Price AL, Reich D. Population structure and eigenanalysis. *PLOS Genetics* 2006; **2**: e190.

15 Price AL, Patterson NJ, Plenge RM, Weinblatt ME, Shadick NA, Reich D. Principal components analysis corrects for stratification in genome-wide association studies. *Nat Genet* 2006; **38**: 904–909.

16 Loh P-R, Danecek P, Palamara PF, Fuchsberger C, A Reshef Y, K Finucane H *et al.* Reference-based phasing using the Haplotype Reference Consortium panel. *Nat Genet* 2016; **48**: 1443–1448.

17 Durbin R. Efficient haplotype matching and storage using the positional Burrows–Wheeler transform (PBWT). *Bioinformatics* 2014; **30**: 1266–1272.

18 Vilhjalmsson BJ, Yang J, Finucane HK, Gusev A, Lindstrom S, Ripke S *et al.* Modeling Linkage Disequilibrium Increases Accuracy of Polygenic Risk Scores. *Am J Hum Genet* 2015; **97**: 576–592.

19 Vattikuti S, Guo J, Chow CC. Heritability and Genetic Correlations Explained by Common SNPs for Metabolic Syndrome Traits. *PLOS Genetics* 2012; **8**: e1002637.

20 Kaiser HF. The Application of Electronic Computers to Factor Analysis. *Educational and Psychological Measurement* 1960; **20**: 141–151.

21 Jackson DA. Stopping Rules in Principal Components Analysis: A Comparison of Heuristical and Statistical Approaches. *Ecology* 1993; **74**: 2204–2214.

22 O'connor BP. SPSS and SAS programs for determining the number of components using parallel analysis and Velicer's MAP test. *Behavior Research Methods, Instruments, & Computers* 2000; **32**: 396–402.

23 Cattell RB. The Scree Test For The Number Of Factors. *Multivariate Behav Res* 1966; **1**: 245–276.
